# On bridging paired-end RNA-seq data

**DOI:** 10.1101/2021.02.26.433113

**Authors:** Xiang Li, Qian Shi, Mingfu Shao

## Abstract

**Motivation:** The widely-used high-throughput RNA-sequencing technologies (RNA-seq) usually produce paired-end reads. We explore if full fragments can be computationally reconstructed from the sequenced two ends—a problem here we refer to as *bridging*. Solving this problem provides longer, more informative RNA-seq reads, and hence benefits downstream RNA-seq analysis such as transcriptome assembly and expression quantification. However, bridging is a challenging and complicated task owing to alternative splicing, transcript noises, and sequencing errors. It remains unclear if the data itself provides sufficient information for accurate bridging, let alone proper models and efficient algorithms that characterize and determine the true bridges.

**Algorithmic Results:** We studied this problem in two settings: reference-based bridging, which assumes reads alignments are available and reconstructs the alignments of full fragments, and *de novo* bridging, which reconstructs sequences of entire fragments from sequences of the two ends. We proposed a novel mathematical formulation that works for both settings—to seek a path in an underlying graph data structure (i.e., splice graph for reference-based bridging, and compacted de Bruijn graph for de novo bridging) such that its bottleneck weight is maximized. This formulation characterizes true bridges and is efficient in filtering out false bridges. This formulation admits optimal substructure property, and hence efficient dynamic programming algorithms can be designed. For reference-based bridging, we designed such an algorithm to calculate the top *N* bridging paths, followed by a voting approach to select one using the distribution of fragment length. For *de novo* bridging, we designed a new truncated Dijkstra’s algorithm. To further speed up, we proposed a novel algorithm that reuses the shortest path tree to avoid running the truncated Dijkstra’s algorithm from scratch for all vertices. These innovations result in scalable algorithms that can bridge all paired-end reads in a compacted de Bruijn graph with millions of vertices.

**Experimental Results:** We showed that paired-end RNA-seq reads can be accurately bridged to a large extend. Our reference-based bridging tool could correctly bridge more than 79.6% of reads. For *de novo* bridging, high precision was observed with varied sensitivity. We also showed that bridging can improve reference-based transcript assembly: the improvement was significant (up to 14.4% measured with adjusted precision), and universal in all combinations with different aligners and assemblers.

**Availability:** Implementations of the algorithms for reference-based and *de novo* bridging are available at https://github.com/Shao-Group/rnabridge-align and https://github.com/Shao-Group/rnabridge-denovo, respectively. Scripts, datasets, and documentations that can reproduce the experimental results in this manuscript are available at https://github.com/Shao-Group/rnabridge-test.

## 1 Introduction

The established high-throughput RNA sequencing technologies (RNA-seq) enables global and accurate measurement of isoform-level gene activities. The second generation short-reads RNA-seq, which remains *de facto* standards for most expression studies, produces paired-end reads. Such data reports sequences of the two ends of a fragment of a RNA molecule, but misses the middle portion of the fragment. The fact that the two ends are from the same molecule and that the length of fragments follows a certain distribution (through fragment size selection) provides valuable long-range information in determining complicated splicing variants, and has been incorporated into various RNA-seq analysis tools and software to improve accuracy, including splicing-aware alignment (e.g., STAR [4], HISAT2 [10], SpliceMap [1]), expression quantification (e.g., Salmon [16], kallisto [2], RSEM [11]), transcript assembly (e.g., StringTie [18], TransComb [13], Scallop [19]), fusion detection (e.g., FuSeq [20], STAR-Fusion [6], SQUID [15]), and alternative splicing quantification (e.g., DARTS [21], leafCutter [12]), among many others.

We explore computationally inferring full fragments from given paired-end RNA-seq reads, a problem we refer to as *bridging*. We study this problem in two settings, namely reference-based bridging and de novo bridging. For reference-based bridging, we assume that the alignments of the paired-end reads are available (which can be obtained using RNA-seq aligners such as STAR and HISAT2), and we aim to reconstruct the alignments of entire fragments. For *de novo* bridging, we aim to reconstruct the sequences of the full fragments from the sequences of paired-end reads.

We believe solving this problem could benefit various downstream RNA-seq analysis such as transcript assembly, isoform-level expression quantification, and splicing quantification. The inferred full fragments likely contain more splicing junctions than individual reads, and hence provide additional long-range information that helps resolve more complicated splicing variants in transcript assembly. Longer sequences will be less ambiguously located to transcripts expressed from the same gene or homologous genes, and hence improves isoform quantification. The reconstructed full fragments may reveal missing junctions in the unsequenced portion, which may lead to more accurate estimation of junction abundance, and hence improves splicing quantification.

However, bridging is a challenging task. It remains unclear if the reads themselves contain sufficient information to enable accurate bridging. Due to the complicated mechanism of alternative splicing and the dynamic nature of transcription and mRNA degradation, different types of splicing junctions will be captured by RNA-seq reads. According to our experiments, in many gene loci, thousands of different splicing junctions can be observed (a majority of them are with low abundance though). For those loci the possible ways of bridging a pair of reads will be enormous, and it’s unclear what signal can be used to determine the true bridge. Besides, sequencing errors and alignment errors also produce false bridges, making this task more challenging to solve.

We explore modeling above bridging problem as a graph problem. Alignments or sequencing reads can be organized by a graph data structure, for instances, the *splice graph* for alignments and de Bruijn graph (dBG) or compacted de Bruijn graph (cdBG) for sequencing reads [3]. Naturally, the sequenced two ends of a fragment can be mapped to the graph and then represented as a pair of paths (two lists of vertices or edges) in the graph. Therefore, the bridging task becomes to find a path that connects the two paths. (Our work follows this framework for bridging; see Section 2.) However, the resulting graphs are often erroneous due to transcript noises and sequencing/alignment errors. It remains open that what is a good characterization of the true path in bridging—in other words, what objective function should be in formulating bridging as a graph optimization problem. In *de novo* bridging, the resulting dBG or cdBG may contain millions of vertices. This urges scalable algorithms for *de novo* bridging.

Although many existing RNA-seq analysis tools use paired-end information, limited efforts have been made to directly reconstruct full fragments. Existing method for reference-based bridging includes MapPER [8]. MapPER implemented a probabilistic framework: starting from splicing junctions of all end reads, MapPER constructed potential splicing paths connecting paired-end reads; an expectation maximization method then assigned likelihood values to all splice junctions and assigned the most probable alignment for each fragment. (However, we were not able to compare with MapPER; see Appendix.) To our best knowledge, no available tool for *de novo* RNA-seq bridging exists.

## 2 Algorithms

We formulate bridging as a new optimization problem (Section 2.1). We then design efficient algorithms for reference-based bridging and *de novo* bridging in Section 2.2 and Section 2.3 respectively.

### 2.1 Formulation

In either setting, the given RNA-seq data can be organized by a graph data structure, i.e., splice graph for reads alignments and compacted de Bruijn graph (cdBG) for sequencing reads (see Sections 2.2 and 2.3 for their constructions). Let *G* = (*V, E*) be the underlying graph. Let *f* be a fragment, for which we know its two sequenced ends. Each end of *f* can be represented as a path in *G*, and *f* can then be represented as a pair of paths in *G*. Note that multiple fragments may correspond to the same pair of paths in *G*; we cluster them into *equivalent classes*. Formally, an equivalent class is a pair of paths (*p*_1_, *p*_2_) in *G* that represent all fragments with two ends being corresponding to p_1_ and *p_2_* respectively.

Let *F* = (*p*_1_ = (*a*_1_, *a*_2_,…, *a_m_*), *p*_2_ = (*b*_1_, *b*_2_,…, *b_n_*)) be an equivalent class, where path *p*_1_ is represented as a list of vertices (*a*_1_,…, *a_m_*) in the graph (the same for *p*_2_). The problem of bridging fragments in *F* becomes to find a path in *G* from *a_m_* to *b*_1_; we call such path as a *bridging path*. We assume that, all fragments in an equivalent class have the same true bridging path. This is because fragments in an equivalent class are similar, as their two ends are mapped to exactly the same list of vertices in the graph. Algorithmically, this assumption allows us to reduce computational load, as all fragments can be bridged in a single run. In the case of reference-based bridging (see Section 2.2), this also enables a robust voting approach to select the best bridging path from a set of candidates by integrating the distribution of fragment length.

We explore what’s a good formulation to find the best bridging path. The main signal we have is the abundances (i.e., the numbers of reads that support vertices/edges of the graph, modeled as weights of vertices/edges; see Sections 2.2 and 2.3). Intuitively, a bridging path supported by more reads are more likely to be the true bridge. We determined that, a formulation that seeks a bridging path with maximized *bottleneck* weight is appropriate for both reference-based bridging and *de novo* bridging, and performs well on experimental comparisons (see Results). Below we describe this formulation.

We define a full ordering of all bridging paths w.r.t. an equivalent class *F* = (*p*_1_ = (*a*_1_, *a*_2_,…, *a_m_*), *p*_2_ = (*b*_1_,*b*_2_,…,*b_n_*)). Let *q*_1_ and *q*_2_ be two arbitrary paths from *a_m_* to *b*_1_ in *G*. Let 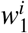 (resp. 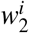) be the ith smallest weight in path *q*_1_ (resp. *q*_2_). We say *q*_1_ is *more reliable* than *q*_2_, if there exists an integer *k* such that 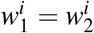 for all 1 ≤ *i* < *k*, and 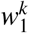 > 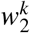. We now formulate the bridging problem as to find the most reliable path. Intuitively, we seek a path *q* from *a_m_* to *b*_1_ in *G* such that the smallest weight in this path is maximized, and in case there are multiple paths with maximized smallest weight, among them we seek the one whose second smallest weight is maximized, and so on.

We believe this formulation is appropriate for bridging paired-end RNA-seq reads. First, by maximizing bottleneck weight, the weakest point of the bridging path is supported strongest, which avoids being dominated by vertices/edges with large weights. Second, through this formulation, false paths coming from sequencing/alignment errors can be efficiently excluded, as they usually exhibits low bottleneck abundance. We also explored formulations such as maximizing the *sum* of weights in a path but they didn’t perform as well as this one.

We emphasize that this formulation satisfies the *optimal substructure* property. Specifically, if *a_m_* → *v*_*i*1_ → *v*_*i*2_ →… → *v*_*i*k_ → *b*_1_ is the most reliable path, then *a_m_* → *v*_*i*1_ → *v*_*i*2_ →… *v*_*i*k_ is the most reliable path from *a_m_* to *v_ik_*. This implies that polynomial-time dynamic programming algorithms can be designed to find the optimal solution.

We use this formulation for both reference-based bridging and *de novo* bridging. For reference-based bridging, instead of just picking the most reliable path, we design an efficient dynamic programming algorithm that finds top *N* paths, and then incorporate the fragment-length distribution to select the best one. For *de novo* bridging, we propose novel, scalable algorithms to identify the most reliable path in large cdBGs with millions of vertices.

The bottleneck weight (smallest weight of the optimal path) in this formulation can be used as a filtering criterion to decide if a bridging path is true. We set this threshold as 1 (i.e., don’t filter) in reference-based bridging but explicitly use this parameter to balance sensitivity and precision in *de novo* bridging (see Table 3 and Table 4).

### 2.2 Reference-based Bridging

#### 2.2.1 Constructing Splice Graph

The input for reference-based bridging is standard RNA-seq alignment in sam/bam format. We first group fragments into gene loci: reads that overlap on their alignment coordinates will be assigned to the same gene locus. We make sure that the two ends from the same fragment will be assigned to the same gene locus. Each gene locus will be treated as independent instances and solved independently.

In each gene locus, junctions will be extracted from spliced reads. The collected set of splicing coordinates will be used to partition the reference genome into continuous segments, called *partial exons*. We use the well-known *splice graph* to represent the partial exons and junctions in a single gene locus (see Figure 1): each partial exon corresponds to a vertex, and each junction corresponds to a direct edge that connects its two corresponding exons. Two additional vertices, source *s* and sink *t*, are added and connected to possible starting and ending partial exons. The weight of an edge is calculated as the number of reads that contain the corresponding junction; the weight of a vertex is calculated as the average coverage of the corresponding partial exon.

**Figure 1:**
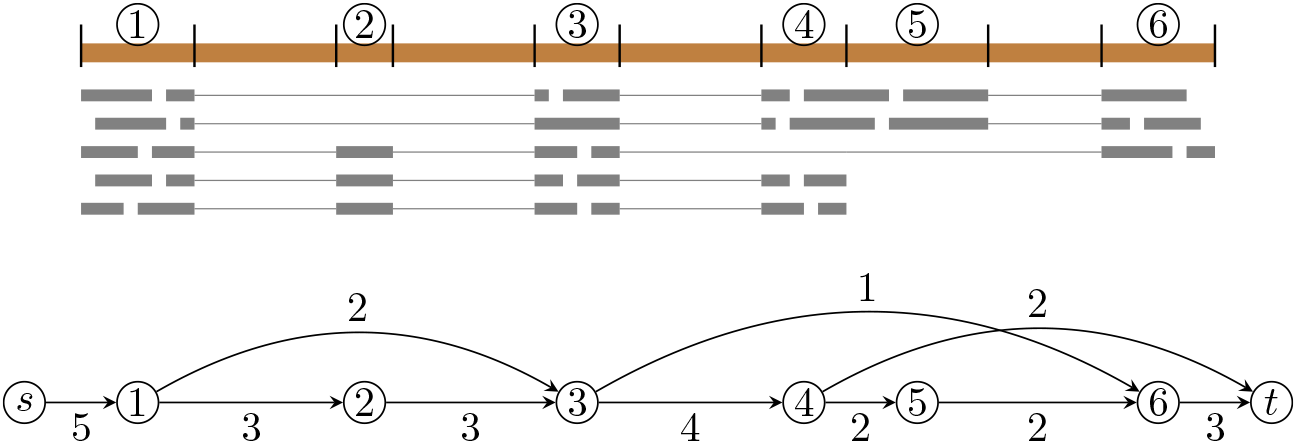
Example of the reads alignment in a gene locus (upper part). Reference genome are portioned into partial exons (numbered from 1 to 6) using the splicing coordinates. Lower part shows the corresponding splice graph; the weight of each edge is given close to it.

#### 2.2.2 Bridging Algorithm

Our core algorithm to bridge all fragments in an equivalent class *F* = (*p*_1_ = (*a*_1_, *a*_2_,…, *a_m_*), *p*_2_ = (*b*_1_, *b*_2_,…, *b_n_*)) consists of two steps: nominating and voting. The nominating step computes the *N* most reliable paths from *a_m_* to *b*_1_ in *G* (in our implementation, *N* is a parameter with default value of 10), and the voting step selects one through a consensus approach using the fragment length distribution.

We implemented a dynamic programming algorithm for nominating. Specifically, let (*v*_1_, *v*_2_,…, *v_n_*) be a topological sorting of all vertices of the splice graph *G* (this is possible because splice graph is a directed acyclic graph). Given a particular *v_i_*, we can use a single run to find *N* most reliable paths from *v_i_* to *v_j_* for every *j* > *i*. To compute these *N* paths for *v_j_*, we examine all vertices *v_k_* that directly connects to *v_j_*, and compare all paths stored in these vertices (each *v_k_* already stores *N* most reliable paths from *v_i_* to *v_k_* at this time point). The best *N* of them will be kept and after concatenating *v_j_* they become the *N* most reliable paths from *v_i_* to *v_j_*. We run this subroutine for all *v_i_*, 1 ≤ *i* ≤ *n*, which gives *N* most reliable paths for all pairs of vertices in *G*. The overall running time of this algorithm is *O*(*N* · |*V* |^2^ · |*E*|).

Above algorithm guarantees finding the most reliable path. To scale better on giant instances with tens of thousands of vertices, we designed a heuristic variant. Instead of maintaining the full list of weights for each path, whose length is *O*(|*V*|), we only store the smallest *M* edge weights (*M* is another parameter with default value of 5) for each path. This gives an improved time complexity of *O*(*N* ·*M* · |*V*| · |*E*|). Although optimality may not be guaranteed, experimental studies show that this heuristic rarely affects the bridging accuracy.

We now describe the voting step. Let (*q*_1_, *q*_2_,…, *q_N_*) be the *N* candidates for an equivalent class *F*. Assume that these *N* paths are sorted (i.e., *q_i_* is more reliable than *q_i+1_*). We now examine each fragment *f* in *F*. For each candidate path *q_i_*, 1 ≤ *i* ≤ *N*, we calculate the actual sequence length of fragment *f*, denoted as | *f_i_*|, assuming that *f* is bridged with *q_i_*. This can be done because once the bridging path of *f* is given, the alignment of the entire fragment *f* is fixed and hence the length of its entire sequence can be calculated. If | *f_i_*| is within a reasonable range (precomputed by sampling 100,000 paired-end reads with unique bridging path, and an empirical distribution of fragment length is then estimated; the value from percentile 2 to percentile 98 will be used as the reasonable range), then fragment *f* votes for path *q_i_*; otherwise, we try next best path (i.e., *q_i+1_*) and check whether |*f_i+1_*| is within the reasonable range. In other words, each fragment votes for the best candidate that leads to a fragment length within the reasonable range. Candidate path with the largest number of votes will be accepted as the bridging path for the equivalent class, and all fragments in it will be bridged using this path. If none of the candidate paths receives any vote, then this equivalent class fails bridging.

#### 2.2.3 Correcting Alignment Errors

We implemented a method to use paired-end information to correct one type of alignment errors while bridging. See Figure 2. Reads that span a junction but flank one side shortly are prone to alignment errors (left red read in Figure 2). Information of paired-end can be used to detect and correct such errors. Let *F* = ((*a*_1_,*a*_2_,…,*a_m_*), (*b*_1_,*b*_2_,…,*b_n_*)) be an equivalent class that fails bridging. If the length of the aligned portion on *a_m_* is small (a parameter with default value of 10 base-pairs), we will then try to bridge *F_1_* = ((*a*_1_,*a*_2_,…,*a*_*m*-1_), (*b*_1_,*b*_2_,…,*b_n_*)). Symmetrically, if *b*_1_ is flanked tinily, we will examine if *F*_2_ = ((*a*_1_, *a*_2_,…, *a_m_*), (*b*_2_, *b*_3_,…, *b_n_*)), or *F*_3_ = ((*a*_1_, *a*_2_,…, *a_m-1_*), (*b*_2_, *b*_3_,…, *b_n_*)) in case both qualify, can be successfully bridged. If so, the corrected and bridged alignments will be reported. In experimental studies (see Table 1), we show that this method could rescue about 1.6-1.9% of the wrongly aligned reads.

**Figure 2:**
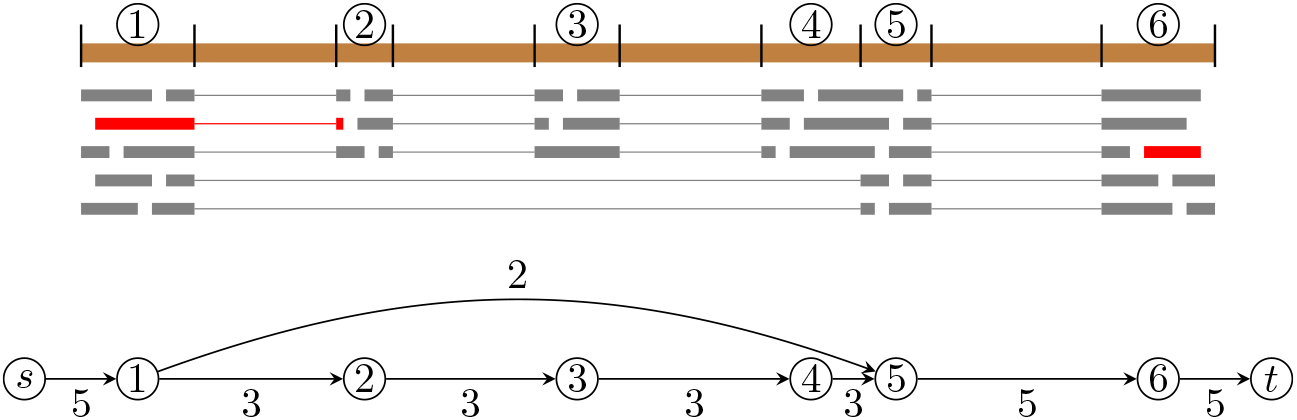
Example of correcting alignments while bridging. One fragment (in red) is in equivalent class ((1,2), (6)). The only bridging path is (2,3,4,5,6), which is most likely beyond the reasonable range of fragment length. As the aligned portion on segment 2 of the first-end is short, we will shorten (1,2) as (1) and then try to bridge it in equivalent class ((1), (6)), for which there is an additional candidate (1,5,6), which likely gives this fragment a sequence length within the reasonable range.

**Table 1:**
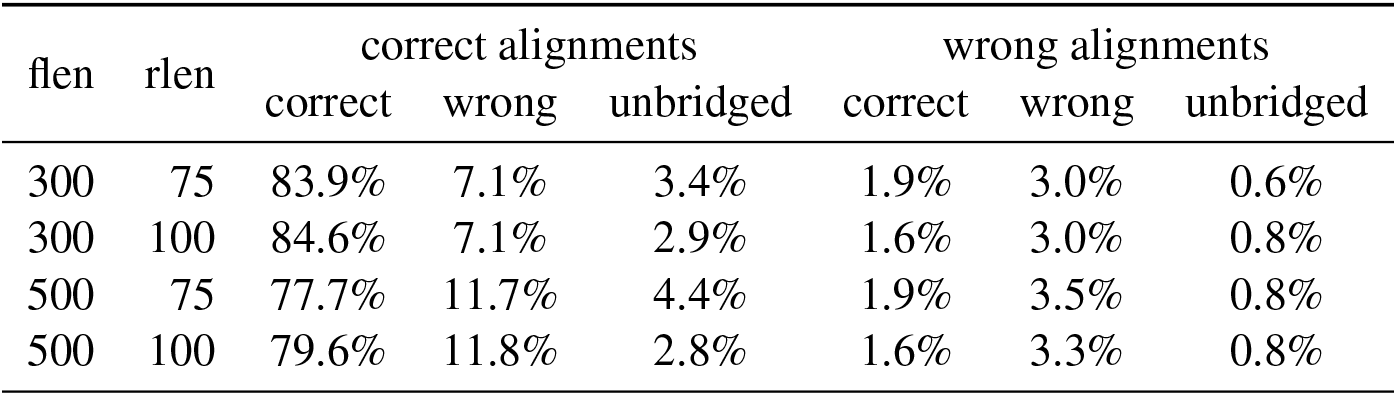
Bridging accuracy on the simulated RNA-seq samples in dataset 1 aligned with STAR. The numbers are normalized by the total number of correctly aligned and wrongly aligned fragments, and averaged over 20 samples for each type. “flen” = average length of fragments; “rlen” = read length.

### 2.3 De Novo Bridging

#### 2.3.1 Constructing Compacted de Bruijn Graph (cdBG)

We use cdBG to represent the given paired-end RNA-seq reads. See Figure 3. In the de Bruijn graph (dBG), each vertex represents a distinct *k*-mer, and its weight equals to the number of appearance this *k*-mer in the reads. The corresponding cdBG is defined as concatenating each simple path (i.e., every vertex in it except the first and the last one has in-degree of 1 and out-degree of 1) of the dBG as a single vertex (the resulting sequence is called a *unitig*). To comply with our formulation of finding the most reliable path, we assign the weight of each vertex in cdBG as the smallest weight of the corresponding simple path in dBG.

**Figure 3:**
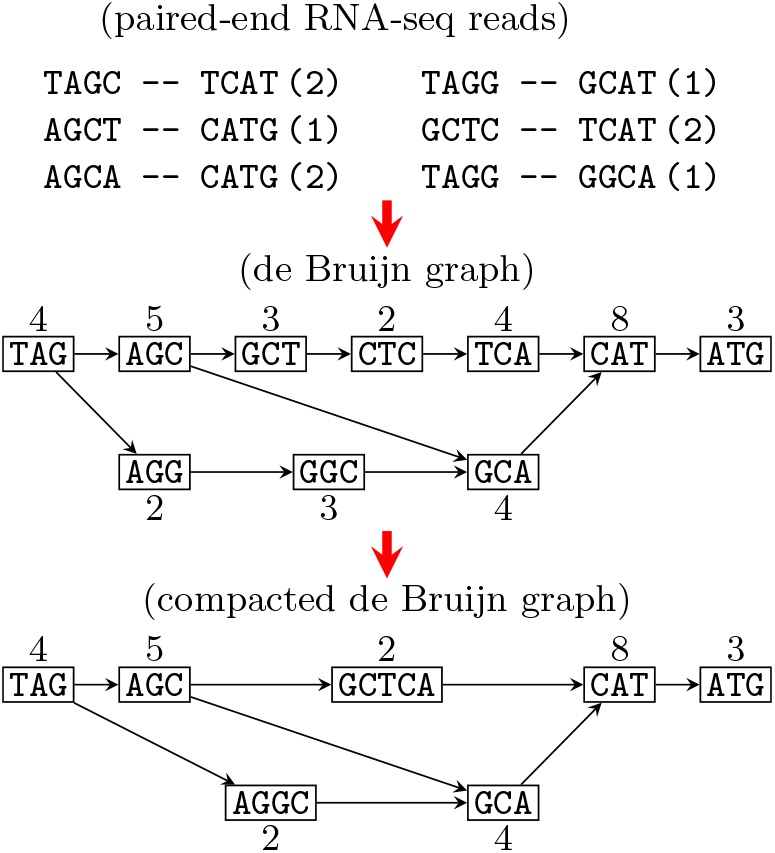
Example of a cdBG constructed from paired-end RNA-seq reads.

In implementation, we directly construct the cdBG and calculate vertex weights, instead of explicitly constructing dBG as an intermediate. Specifically, we use library Bifrost [7] to build the cdBG. In order to assign vertex weights, we first build a hashing table that stores the frequency of each *k*-mer; we then examine all k-mers in this vertex (a unitig of length *l* contains *l* − *k* + 1 *k*-mers) and assign the smallest frequency as the weight of the vertex.

#### 2.3.2 Bridging Algorithms

Following our formulation, we aim to find the most reliable bridging path for all equivalent classes. As the cdBG constructed from a typical RNA-seq sample may consist of millions vertices (see Table 5), it is simply not affordable to run a standard dynamic programming algorithm for every possible starting vertices (note that a single run of dynamic programming algorithm starting from a vertex will find the most reliable path connecting it to all other vertices). We propose two algorithmic innovations to address this main challenge in *de novo* bridging. First, we implemented a *truncated Dijkstra’s algorithm* to find the most reliable bridging path starting from a starting vertex *a_m_* to any other vertex up to a certain length *D* (default value is 400 + 2L where *L* is read length). This is to incorporate the prior knowledge that fragment length follows an empirical distribution and an upper bound of fragment length can be assumed. In our truncated Dijkstra’s algorithm, for each vertex *v* we maintain the total length of the most reliable path from the current starting *a_m_* to *v*, and when such length for *v* reaches D, we won’t extend *v* in the Dijkstra’s algorithm. Similar to the algorithm in reference-based bridging, a list that stores the smallest *M* weights also needs to be maintained.

The second algorithmic innovation is that we reuse the optimal solutions obtained for the current starting vertex *a_m_* to construct the optimal solutions for next starting vertex 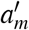. This allows us to avoid running above truncated Dijkstra’s algorithm from scratch for all possible starting vertices. Specifically, for the current starting vertex *a_m_*, we maintain a *shortest path tree*, denoted as *T*(*a_m_*), to store the most reliable paths from *a_m_* to all other vertices (up to length *D*), i.e., the unique path in *T* from root *a_m_* to any vertex *v* gives the most reliable path from *am* to *v* in the cdBG. See Figure 4. This tree can be constructed in linear time with a tracing back procedure after running the truncated Dijkstra’s algorithm. We note that, again according to the property of optimal substructures, any subtree of *T*(*a_m_*), say the one rooted at *u*, also gives the most reliable path from *u* to any other vertex *w* in the subtree. This suggests we can reuse the subtree rooted at *u* to calculate all reliable paths starting from *u* (and then construct the corresponding shortest path tree). More specifically, in implementation we directly load the subtree rooted at *u* to the priority queue (recall that the core data structure of Dijkstra’s algorithm is a priority queue that gets updated iteratively). In other words, for next starting vertex *u* we run the truncated Dijkstra’s algorithm in the middle rather than from scratch, as we already know the optimal solutions for a subset of vertices (i.e., those in the subtree of *T*(*a_m_*) rooted at *u*). To benefit from this property to the largest extent, we determine the starting vertex whose subtree is largest as the next one to bridge.

**Figure 4:**
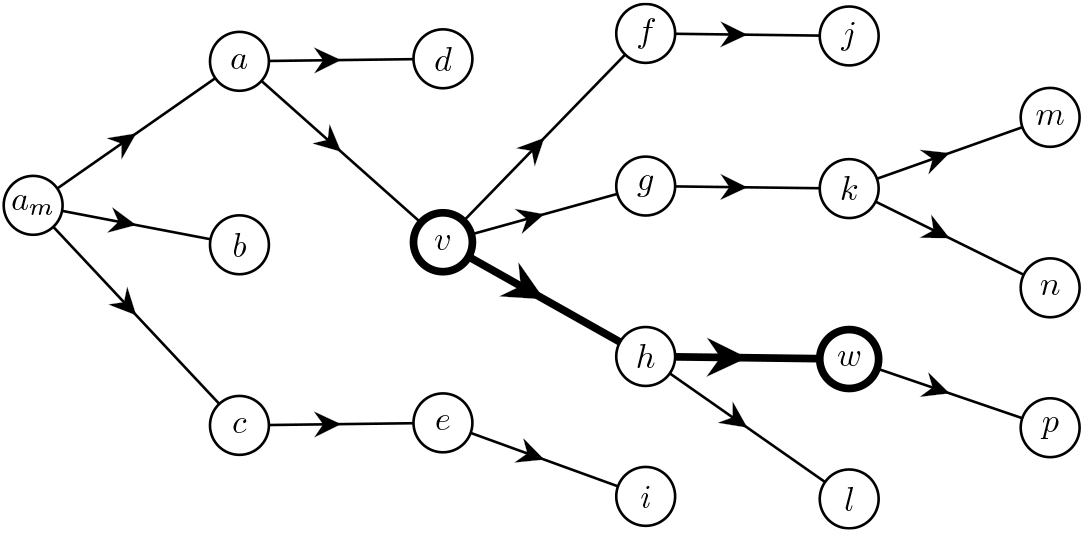
Illustrating the shortest path tree starting from vertex *a_m_*. Consider subtree rooted at *v*. Then for any vertex in it, say *w*, the path from *v* to *w* in this subtree is the most reliable path from *v* to *w* in the cdBG.

## 3 Results

### 3.1 Resulting Tools

We developed a tool for reference-based bridging, available at https://github.com/Shao-Group/rnabridge-align. The input for this tool is the alignments of paired-end reads, in standard sam/bam format. The output is the alignments of entire fragments, again in sam/bam format. The algorithms for *de novo* bridging were implemented as another tool, available at https://github.com/Shao-Group/rnabridge-denovo. The input for it is sequencing reads in fasta format, and it generates sequences of full fragments again in fasta format.

### 3.2 Datasets

We use 2 datasets to evaluate the accuracy of bridging and its effectiveness in improving downstream RNA-seq analysis. The first dataset includes 80 paired-end RNA-seq samples simulated using Flux-Simulator [5]. We vary two parameters in the simulation: the average length of fragments (300 and 500 were used) and the read length (75 and 100 were used). For each combination of parameters, we independently simulated 20 samples. The number of reads in samples with fragment length being 300 and 500 are roughly 150M and 90M, respectively. The second dataset was previously used in the Scallop paper [19]: it contains 10 biological RNA-seq samples.

### 3.3 Accuracy of Reference-based Bridging

We evaluate the bridging accuracy of rnabridge-align using simulation data, for which the ground-truth (i.e., full fragments) are available. We define a bridged fragment is correct only if it’s exactly the same as the ground-truth fragment. As the input for rnabridge-align is the alignments produced by another aligner (we use STAR here), which may not be perfect, we evaluate the bridging accuracy separately on the fragments that are correctly and wrongly aligned.

The averaged accuracies for each simulation type are summarized in Table 1. Overall the bridging accuracy is high with 85.8-86.2% being correctly bridged when the average fragment length is 300. For fragment length of 500 the accuracy slightly drops to 79.6-81.2%. This is expected as longer fragments are harder to bridge and the coverage of these samples are lower. Notice that rnabridge-align also correctly bridges 1.6-1.9% fragments even when the paired-end alignments are wrong, thanks to the procedure in Section 2.2.3.

### 3.4 Improving Reference-based Transcript Assembly

Reference-based bridging infers the alignment of full fragments, which provides longer alignments with enhanced long-range information. Such information is particularly useful in assembling complicated splicing variants. Below, we evaluate the effectiveness of reference-based bridging in improving transcript assembly.

#### 3.4.1 Pipeline

We follow the workflow illustrated in Figure 5. For each aligned RNA-seq sample, we compare directly assembling with first bridging using rnabridge-align followed by assembling the bridged alignments. For each route, we test two recent and leading assemblers StringTie (version 2.1.4; released on 7/7/2020) and Scallop (version 0.10.5; released at 5/22/2020).

**Figure 5:**
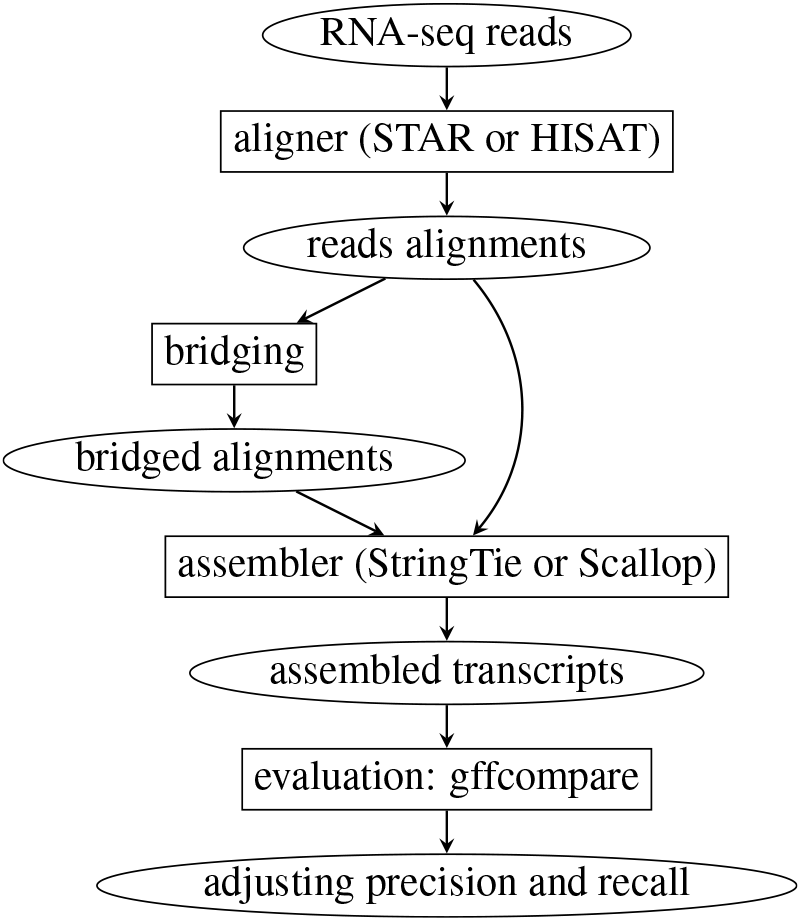
Pipeline of evaluating rnabridge-align in improving transcript assembly.

#### 3.4.2 Evaluation Measures

Each method assembles a set of transcripts. We define an assembled transcript is *known* if its intron-chain coordinates exactly match those of an existing transcript in the reference transcriptome. We use a third-party tool, gffcompare [17], to compute the number of known transcripts (proportional to sensitivity), and precision (the ratio between the number of known transcripts and the total number of predicted transcripts).

Notice that one method might assemble more known transcripts but exhibit lower precision, or vise versa, on certain samples. To enable comparing in these cases, we first balance them in one measure and then compare the other one. Specifically, let *k*_1_ and *k*_2_ be the number of known transcripts and *p*_1_ and *p*_2_ be the precisions obtained by two methods *X*_1_ and *X*_2_. Assume that *k*_1_ > *k*_2_. We sort the assembled transcripts obtained by *X*_1_ based on the expression abundance, and gradually filter out them starting from ones with low abundance. In this way, the known transcripts of *X*_1_ will decrease while its precision will likely increase (as lowly-expressed ones are more likely to be false positive transcripts). We calculate the two measures for *X*_1_, denoted as 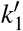 and 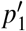, as filtering goes, and stop when 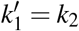. We then report 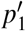 at this time as the *adjusted precision* of *X*_1_. Note that the relationship between 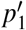 and *p*_2_ can therefore reflect the overall performance of the two methods, as their numbers of known transcripts have been adjusted identical. Similarly, we can compute the *adjusted number of known transcripts*, through gradually filtering out transcripts with lowest abundance in the method with lower precision, until their (adjusted) precisions match. This way of adjusting can be generalized into multiple methods; in our case, we have 4 methods. Such adjusted measures have been used in evaluating assembly accuracy [19].

#### 3.4.3 Results on Simulated Dataset (Dataset 1)

The average assembly accuracies of the 4 methods on dataset 1 are illustrated in Figure 6 and Figure 7. For the category with average fragment length being 300 and read length being 75, the average adjusted precisions for StringTie with and without using bridging are 67.3% and 72.7% respectively, and such numbers for Scallop are 56.2% and 70.6%, respectively; in adjusted known transcripts, StringTie with and without using bridging assembled 6091 and 6421 known multi-exon transcripts respectively (5.4% increase), and such numbers for Scallop are 5575 and 6137 respectively (10.1% increase). Similar or less improvements are noted, but overall bridging improves transcript assembly on all 4 types of simulations using any of the two assemblers.

**Figure 6:**
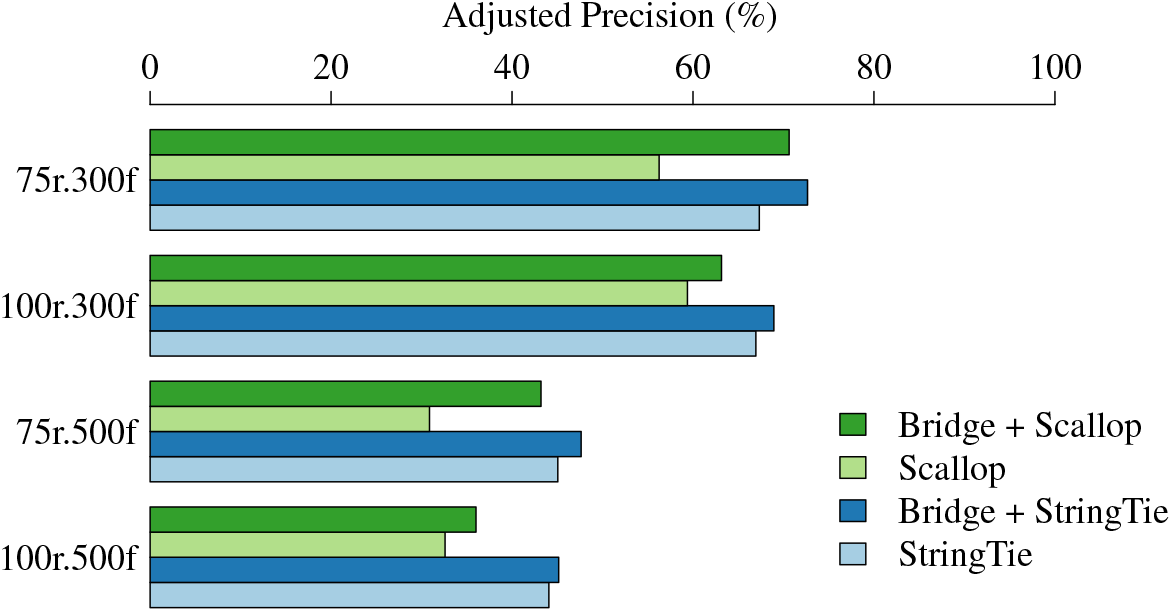
Comparison of the average adjusted precision on 4 types of simulated RNA-seq data. All samples are aligned with STAR.

**Figure 7:**
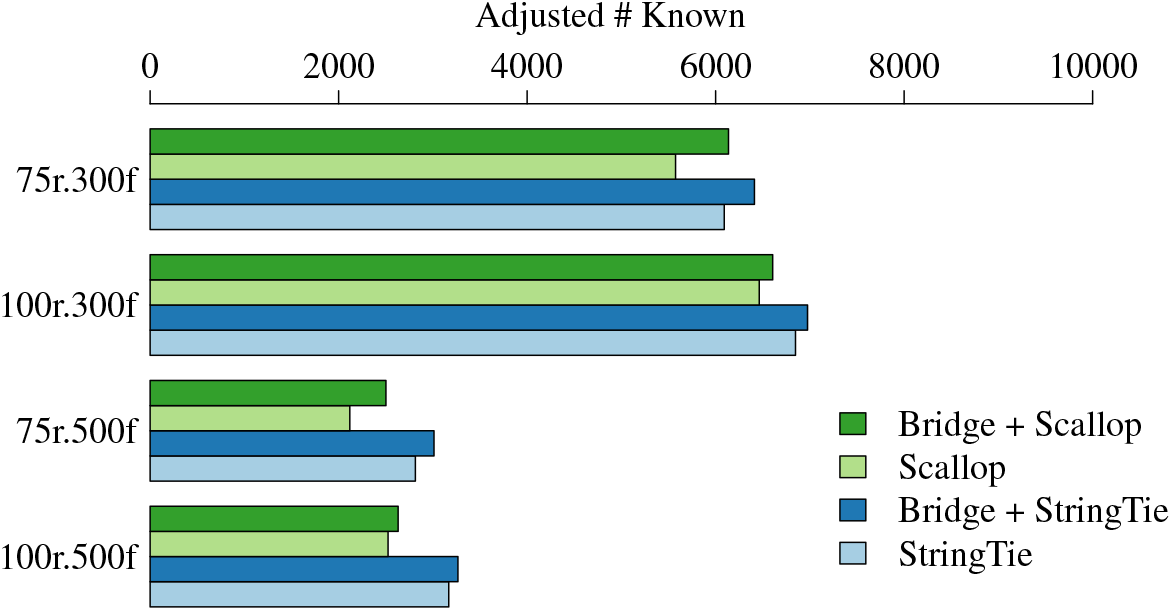
Comparison of the average adjusted number of known transcripts on 4 types of simulated RNA-seq data. All samples are aligned with STAR.

#### 3.4.4 Results on Biological Dataset (Dataset 2)

The average assembly accuracies over the 10 RNA-seq samples in dataset 2 are shown in Figure 8 and Figure 9. Again bridging improves assembly in all settings. For examples, when STAR is used, the average adjusted precisions for StringTie with and without using bridging are 32.9% and 40.6% respectively, and such numbers for Scallop are 44.0% and 46.1%, respectively; in adjusted known transcripts, StringTie with and without using bridging assembled 14344 and 13303 known multi-exon transcripts (7.8% increase), respectively, and such numbers for Scallop are 15590 and 16012 (2.7% increase), respectively. Similar improvements are noted when HISAT2 is used.

**Figure 8:**
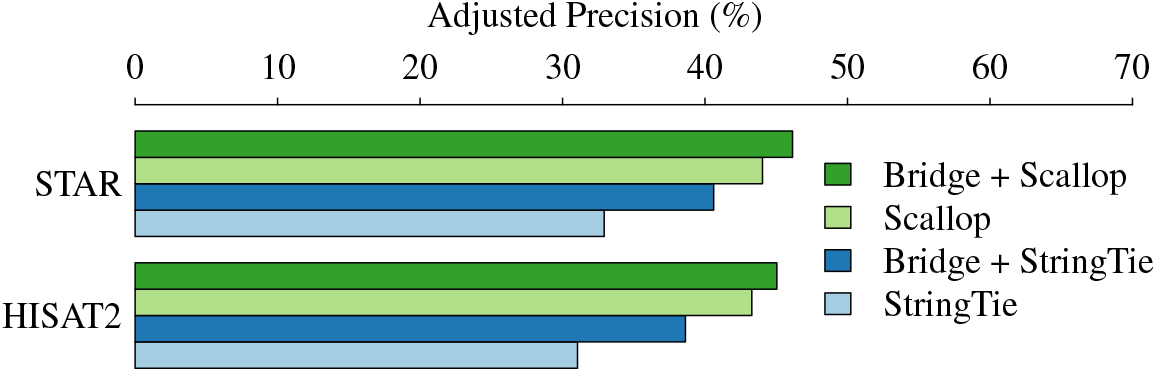
Comparison of the average adjusted precision on dataset 2.

**Figure 9:**
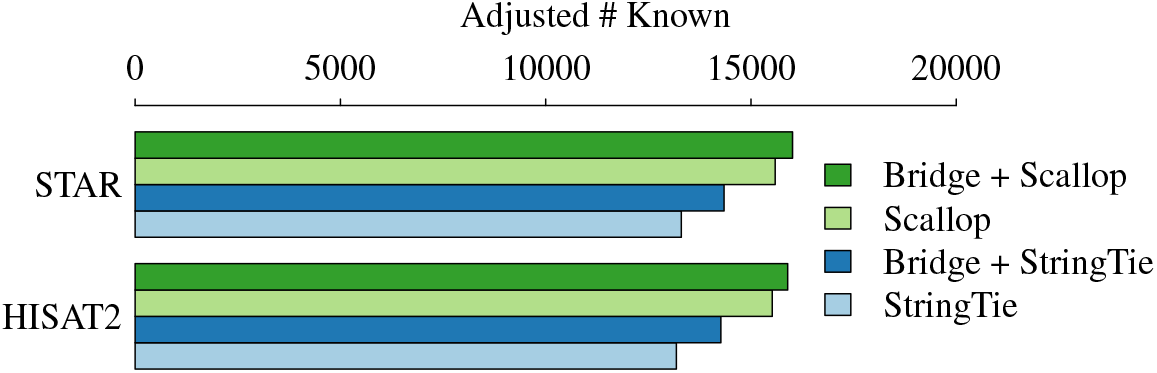
Comparison of the average adjusted number of known transcripts on dataset 2.

#### 3.4.5 Further Analysis

Transcript assembly has long been recognized as a very challenging but critical task. We think the improvement achieved by bridging RNA-seq reads is quite significant: up to 14.4% for adjusted precision and 10.1% for adjusted sensitivity. Although such increase varies, improvements are observed on *all* settings with different aligners, assemblers, and simulation parameters. This confirms that our bridging tool is robust in improving reference-based transcript assembly.

StringTie and Scallop exhibit differently on simulated and biological data. This has also been observed in other studies. One of the possible reasons is that Scallop has been tailored to identify false positives during assembly (in general biological RNA-seq data is more noisy than simulated data). Again, the focus here is to show the improvement when bridging is incorporated, regardless of downstream assembler used.

On biological samples, the adjusted precision (ranging from 30%-50%) is low comparing with simulated samples with fragment length being 300. This is because we use reference transcriptome as ground-truth (as for these biological RNA-seq data we don’t know the true expressed transcripts). The current transcriptome has been regarded as incomplete and over-annotated. Therefore, the precision and known transcripts reported here underestimated the actual performance of all approaches. But these measures are fair for all methods in comparing their *relative* performances. The low precision for simulated samples with fragment length being 500 are likely due to that they are harder instances and that these samples are with low coverage.

#### 3.5 Running Time of Reference-based Bridging

The running time of rnabridge-align on samples in dataset 2 is given in Table 2. This is faster or comparable to assemblers like StringTie or Scallop on most biological RNA-seq samples. Simulated samples are less noisy and rnabridge-align runs even faster on them.

**Table 2:** CPU time (in minutes) of rnabridge-align when running on the 10 samples in dataset 2 aligned with STAR or HISAT2.

| SRA ID | time (STAR) | time (HISAT 2) |
| --- | --- | --- |
| SRR307903 | 8.9 | 7.2 |
| SRR307911 | 8.9 | 8.8 |
| SRR315323 | 7.5 | 6.2 |
| SRR315334 | 9.0 | 7.9 |
| SRR387661 | 25.0 | 24.1 |
| SRR534291 | 23.2 | 23.9 |
| SRR534307 | 32.3 | 33.7 |
| SRR534319 | 15.3 | 15.1 |
| SRR545695 | 25.0 | 25.4 |
| SRR545723 | 9.1 | 8.6 |

#### 3.6 Accuracy of de novo Bridging

We first evaluate the accuracy of rnabridge-denovo using simulation data, for which the ground-truth are available. We define a bridged fragment is correct only if it’s exactly the same as the ground-truth fragment. The sensitivity is defined as the number of correctly bridged fragments divided by the total number of fragments (i.e., paired-end reads), and precision is defined as the number of correctly bridged fragments divided by the total number of bridged fragments.

The results are summarized in Table 3. The bottleneck threshold used to filter bridges is set to 5 for all simulated samples. Overall our tool exhibits high accuracy. The accuracy drops with long fragment length. This is expected as in this case the missing portion is longer and therefore harder to bridge. This may also be partly due to that the coverage of these simulated samples is low. Higher accuracy is observed when the read length is longer at the same fragment length, again as expected.

We then evaluate using the biological data in dataset 2. As we don’t have ground-truth for them, we again use the sequences in the reference transcriptome to evaluate. We align all the bridged fragments to reference using BLAT [9]. We define a bridged fragment is correct only if it is hit by one of the reference sequences with at least 95% sequence identity. The sensitivity and precision is defined the same as in evaluating with simulation data.

**Table 3:** Averaged accuracy of rnabridge-denovo on the simulated RNA-seq samples in dataset 1.

| flen | rlen | bottleneck | sensitivity (%) | precision (%) |
| --- | --- | --- | --- | --- |
| 300 | 75 | 5 | 80.7 | 91.8 |
| 300 | 100 | 5 | 85.7 | 92.5 |
| 500 | 75 | 5 | 66.6 | 85.4 |
| 500 | 100 | 5 | 76.3 | 86.9 |

The results are summarized in Table 4. Overall the precision keeps high but the sensitivity varies quite a lot. This is because we prioritize precision by setting a high bottleneck-threshold: 20 for samples with less than 50M paired-end reads and 100 otherwise. Comparing with simulated data, biological data are more noisy and harder to bridge.

**Table 4:** The accuracy of rnabridge-denovo on the 10 RNA-seq samples in dataset 2. The number of paired-end reads are given in unit of million.

| SRA ID | #paired-reads | bottleneck | sensitivity (%) | precision (%) |
| --- | --- | --- | --- | --- |
| SRR307903 | 36.0M | 20 | 65.3 | 91.8 |
| SRR307911 | 41.4M | 20 | 42.7 | 91.3 |
| SRR315323 | 30.3M | 20 | 27.9 | 87.9 |
| SRR315334 | 39.5M | 20 | 49.5 | 93.3 |
| SRR545723 | 38.9M | 20 | 42.7 | 81.6 |
| SRR387661 | 124M | 100 | 58.2 | 90.1 |
| SRR534291 | 114M | 100 | 64.3 | 93.5 |
| SRR534307 | 165M | 100 | 50.6 | 74.7 |
| SRR534319 | 76.6M | 100 | 24.6 | 69.5 |
| SRR545695 | 119M | 100 | 37.6 | 75.5 |

#### 3.7 Running Time of de novo Bridging

Our tool rnabridge-denovo consists of 3 modules. Module 1 uses Bifrost [7] to build the cdBG; module 2 aligns all paired-end reads to the cdBG and calculates the weight of unitigs; module 3 implements our core bridging algorithm described in Section 2.3.2. We show the breakdown of the CPU time of the 3 modules together with the size of the resulting cdBGs on the 10 biological samples in Table 5. Note that module 3 takes the least CPU time among 3 modules, which proves the efficiency and scalability of the optimized core bridging algorithms.

**Table 5:** The size (number of vertices and edges; in unit of million) of the cdBGs and the CPU time (in minutes) measured for the 3 modules of rnabridge-denovo running on the 10 samples in dataset 2.

| SRA ID | size of cdBG |  | CPU time (in minutes) |  |  |
| --- | --- | --- | --- | --- | --- |
|  | #vertices | #edges | module 1 | module 2 | module 3 |
| SRR307903 | 5.15M | 6.99M | 149 | 57 | 26 |
| SRR307911 | 8.11M | 10.71M | 175 | 66 | 75 |
| SRR315323 | 7.68M | 9.45M | 120 | 40 | 63 |
| SRR315334 | 6.35M | 10.16M | 158 | 67 | 37 |
| SRR387661 | 18.71M | 27.36M | 515 | 223 | 141 |
| SRR534291 | 15.14M | 26.67M | 685 | 370 | 128 |
| SRR534307 | 34.29M | 56.00M | 1038 | 561 | 281 |
| SRR534319 | 16.48M | 22.09M | 311 | 118 | 121 |
| SRR545695 | 19.43M | 27.49M | 479 | 201 | 174 |
| SRR545723 | 13.47M | 16.04M | 243 | 99 | 159 |

## 4 Conclusion

We conclude that the formulation of finding the path with maximum bottleneck weight (breaking ties with second bottleneck weight, and so on) is an appropriate model for both reference-based bridging and *de novo* bridging. This formulation admits optimal substructure, a nice property that leads to polynomial-time, exact algorithms. The new techniques we introduced in *de novo* bridging, in particularly reusing shortest path tree to speed up, are likely of independent algorithmic interests in other applications involved in cdBGs.

On both simulated and biological data, the precision of bridged reads is high (around 80-90%), suggesting that our tools can be safely combined with other RNA-seq tools and pipelines. The sensitivity, although not consistent, is quite significant (above 50% for most cases). This suggests a positive answer to the question that whether the data itself contains enough information for bridging.

We also showed that our reference-based bridging tool could improve reference-based transcript assembly. Such improvement is universal, regardless of involved aligner, assembler, or simulation parameters.

Both tools are modular, with input and output being standard sam/bam/fasta formats. Therefore they can be easily incorporated into existing RNA-seq analysis pipeline. We devote to reproducibility: the experimental results in this manuscript can be reproduced with the scripts available at GitHub.

## 5 Discussion

We explored if *de novo* bridging could improve *de novo* transcript assembly. To this end we piped the bridged fragments to one leading assembler TransLiG [14], but only observed marginal improvement. This may be because TransLiG is not optimized to make use of mixed short and long sequences. (On the contrary, some reference-based assemblers like Scallop and StringTie does take advantage of reads that thread multiple vertices to improve assembly accuracy, and hence benefit from bridged fragments.) Developing a new *de novo* assembler that can fully use such bridged data is on our research agenda. Experimenting if *de novo* bridging could improve isoform quantification and splicing quantification is also an interesting future research topic for us.

The sensitivity of rnabridge-denovo is low on some biological samples. One reason is that we use a high bottleneck-threshold to keep high precision, which consequently disconnects paired-end reads in low-coverage gene loci. We are developing a post-bridging algorithm to use full-range information in the reads to decide if a bridged fragment is correct (rather than just using a bottleneck-threshold), in a hope of keeping high precision while improving sensitivity. Specifically, note that the cdBG is not a loss-free representation of sequencing reads, as it breaks reads into *k*-mers, and any phasing information beyond (*k* + 1)-mer is not represented. Let *s* be the bridged sequence of a fragment *f* constructed using above algorithm. We can examine each sliding window of length *L*, where *L* is the read length, and determine the number of input reads that are identical to this L-mer (i.e., these reads *support* this L-mer). This gives a *supporting profile*, a vector of length |*s*| — *L* + 1 for this bridged sequence. Intuitively, a profile with few 0s suggests that the bridged sequence is likely a true one, while a long consecutive 0s suggests a false bridge. We are experimenting if such supporting profile could lead to more efficient algorithms in boosting bridging accuracy.

## Appendix

We were not able to compare with MapPER in our experiments. MapPER requires input sam file to be in a special format (referred to as “PER_SAM”). We looked into MapPER’s source code and it seems that MapPER only deals with CIGAR fields with “M” (alignment match) and “N” (skipped region), but for biological samples there are plenty of “S” (soft clipping). On simulated data we observed that MapPER generated “*” for the CIGAR field in many alignments. We contacted the authors of MapPER but haven’t get responded.

